# Using Genome Wide Estimates of Heritability to Examine the Relevance of Gene-Environment Interplay

**DOI:** 10.1101/037861

**Authors:** Ben Domingue, Jason Boardmani

## Abstract

We use genome-wide data from the third generation respondents of the Framingham Heart Study to estimate heritability in body mass index using different quantities of the measured genotype. Heritability decreases rapidly when SNPs implicated by a genome-wide association study are removed but shows essentially no decline when SNPs implicated by a gene-environment interaction in a second genome-wide analysis are removed. This second result is highlighted by our additional finding that the SNPs which explain heritability amongst a subsample defined by higher educational attainment explain no heritability of the heritability in the lower education group, and vice-versa. Finally, we do find consistent heritability estimates when we compare family-based estimates versus those based on measured genotype.

## 1 Introduction

Evidence from twin and sibling studies suggests that most behavioral phenotypes of interest to social demographers evidence moderate to large heritability estimates. Traits such as smoking, drinking, obesity, and exercise are all found to be roughly 40-60% heritable. These large estimates of genetic influence are striking given that, to date, genome-wide association studies (GWAS) have not uncovered SNPs or SNP clusters that explain more than 1-2% of phenotypic variance. The so called “missing heritability” (Maher, 2008; Manolio et al., 2010) has defined the first generation of GWAS studies and has led to a re-thinking of standard approaches.

From a social science perspective, BMI is an interesting phenotype since it has strong biological and social components. There is strong evidence that genes determine individual differences in physical weight and weight gain (Haberstick et al., 2010; Fox et al., 2007; W. Yang, Kelly, & He, 2007). There is also a great deal of variability in the estimated influence of genotype on BMI; with an average of roughly 60%, heritability estimates for BMI range from as little as 5% to as high as 90% (Loos & Bouchard, 2003). This variation is in line with the GxE perspective that anticipates differential associations between genotype and phenotype across different environments (Shanahan & Hofer, 2005) and some work has demonstrated the social moderation of genetic factors linked to obesity related phenotypes (Lee, Lai, Ordovas, & Parnell, 2011; Boardman et al., 2012).

More recent models that use genetic similarity among unrelated persons have begun to provide estimates of genetic influence that are similar to traditional behavior genetic results (J. Yang et al., 2010; J. Yang, Lee, Goddard, & Visscher, 2011). These methods have been organized under the umbrella of GCTA and a toolkit for these analyses is available to all researchers.1 To date, these methods have been primarily used to characterize the relative contribution of additive genetic influences on overall phenotypic variance. No existing study has used these methods to explore the relevance of the gene-environment interaction perspective. The purpose of this study is to use the GCTA methodology to decompose the variation of BMI into components that are due to main and interactive effects. We do this by estimating heritabilities after repeatedly removing SNPs that were identified by various genome-wide analyses as top hits. Our primary finding is that the top hits identified by a genome-wide search for SNPs that differentially predict phenotype by environment account for virtually none of the heritability.

## 2 Methods

The GCTA model (J. Yang et al., 2011) has become a popular alternative to using sibling and twin data to estimate heritability. Rather than using expected genetic similarity, such as of .5 and 1 in the case of dizygotic and monozygotic twins, the GCTA models use the empirical genetic similarity score the characterizes genetic similarity above and beyond what we would expect by chance (e.g., IBS). This N×N genetic relationship matrix is then used to partition variance into components that are due to measured genetic associations or those due to environmental factors.

Our contribution is to focus on the results from removing different blocks of SNPs when calculating GCTA-based heritablities. That is, we remove blocks of SNPs and subsequently recalculate heritability to see how much of the genetic variation is explained by the removed SNPs. The SNPs are removed in specific orders, typically based on a ranking of their associated p-values from GWAS analyses. The first model is a typical GWAS:

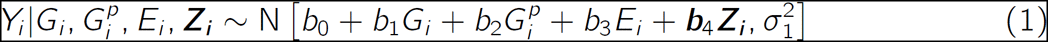

where bold characters represent multidimensional components (either vectors or matrices). Equation 1 describes BMI (*Y*) as a function of genes (*G*) and other factors. First, a control for parental mating type (*G^p^*) is included. Parents may both be homozygous for the minor allele (AA, AA), one may be homozygous for the minor and the other homozygous for the alternate allele (AA, BB), they may both be heterozygous (AB, AB), etc. In total, there are six potential mating types. The use of trios (genetic data for parents and their offspring) enables researchers to look at within family distributions of alleles and this approach is robust to population stratification. Second, a control for environment (E_*i*_), an indicator of whether an individual has obtained a college degree is included. Finally, additional controls for gender and age (**Z**_*i*_) are also included.

We then use a genome-wide gene-environment interaction (GWGEI) model that controls for gene-environment correlation as well as population stratification. This model is proposed by Moreno-Macias, Romieu, London, and Laird (2010) and has been shown to provide valid and reliable GWGEI estimates. The model is:

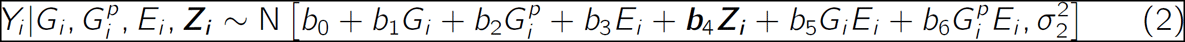

This model differs from the above in the introduction of the interactions. The first interaction (*b*_5_*G_i_E_i_*) allows for the effect of genotype to differ by environment. The second interaction 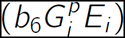 reduces the risk of gene-environment correlation which may have important influences on GxE parameter estimates. This model is comparable to the FBAT approach (Laird, Horvath, & Xu, 2000; Laird & Lange, 2006) that has been successfully used to identify SNPs linked to obesity (Herbert et al. 2006).

The models above emphasize allelic associations and corresponding interactions one SNP at a time. We now consider how this information can be used to adapt existing quantitative genetic modeling techniques focused on genetic similarity among unrelated individuals (J. Yang et al., 2010, 2011). These methods have been organized into the GCTA suite of genome wide association tools. Rather than estimating the cumulative influence of all known causal loci (which are, in principle, unknown), these models estimate a relationship matrix for all unrelated persons. They characterize the genetic relationships between individuals *j* and *k* individual across all genetic markers *i* with minor allele frequency *p_i_* as:

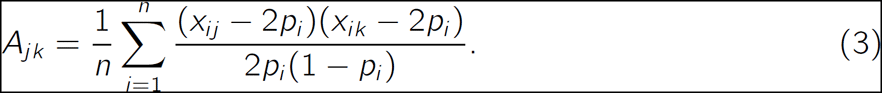

Related pairs will have values that, on average, correspond to the fixed values in traditional behavioral genetic models (e.g., siblings will have an average of .5) but the inclusion of family members can artificially inflate heritability estimates because the shared environment is subsumed in the family coefficient. As such, these researchers recommend eliminating pairs with *A_jk_* estimates in excess of .025. The matrix of genetic similarity estimates is then used in the estimation of:

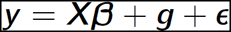

where ***є*** is white-noise error and ***g*** ∼ Normal 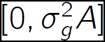. Heritability is then defined as the ratio of the genetic variation to the total variation:

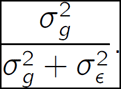

Details on estimation can be found in J. Yang et al. (2011). A critical examination of the performance of this model is beyond the scope of this project, please see Speed, Hemani, Johnson, and Balding (2012); Kumar, Feldman, Rehkopf, and Tuljapurkar (2016); Conley et al. (2014); Domingue et al. (in press).

## 3 Data

The study sample for this project was derived from the Framingham SNP (Single Nucleotide Polymorphism) Health Association Resource (SHARe, version 6) as available through NCBI’s database of Phenotypes and Genotypes (dbGaP). The analysis for this study focused on the third generation of the Framingham Heart Study (FHS, Splansky et al., 2007). The original cohort of the FHS was first assessed in 1948 and nearly 25 years later their children and many of their spouses participated in the offspring cohort study (W2). Then, in 2002, roughly 4000 adults who had at least one parent in the offspring cohort took part in the third generation (W3) cohort. This cohort was examined for a variety of different morbidities using clinical and laboratory assessments. Crucially, study participants were measured for height and weight.

Genetic samples were available via the Framingham SHARe resource which contains genotypes for all respondents using the Affymetrix 5.0 genotyping platform. After reducing the Framingham SHARe data set to trios with complete (non-missing) genetic information (e.g., genotypes for biological mother, biological father, and focal subject) our analytic sample includes 1,967 W3 respondents. We drop SNPs with minor allele frequencies of less than 5% and those that do not meet Hardy Weinberg criteria.^2^ This initial pruning was done in PLINK and 256,884 SNPs met this criteria.

Descriptives for the phenotype and individuals used here are shown in Table 1. The W3 respondents had higher average BMIs (26.6) than the W2 respondents (25.5). Of the W3 respondents, those with a college degree had lower BMIs than those without (26.0 compared to 27.4). This is a moderate effect size and translates to roughly 10 additional pounds for a 140 pound adult who is 5’8". There are no gender differences by educational attainment but we do find that the college educated sample is slightly younger. We include these controls in subsequent analyses. GCTA Heritabilities were also computed based on the full W3 sample as well as when it was split by having a college degree. For the full sample, the heritability of BMI was 0.5 although heritabilities were higher (though not significantly so) in the samples split by education.

**Table 1:** Descriptive statistics for BMI

|  | Mean (SD) | N |
| --- | --- | --- |
| W2 | 25.5 (4.18) | 1624 |
| W3 | 26.6 (5.44) | 1979 |
| W3 College | 26.0 (5.2) | 1103 |
| W3 Non-College | 27.4 (5.6) | 864 |
| | $h^2$ (SE)* | N |
| W3 | 0.50 (0.05) | 1979 |
| W3 College | 0.60 (0.08) | 1103 |
| W3 Non-College | 0.53 (0.09) | 864 |

\* Based on GCTA estimates.

## 4 Results

Figure 1 compares the declines in heritability as SNPs are removed in blocks of 5000 in four different orders. When no SNPs are removed (the point where all four curves touch on the y-axis), heritabilities are all identical. Consider first the line with the most severe negative slope. This line, labeled GWAS, represents the repeated removal of the SNPs that had the lowest p-values from the GWAS analysis (p-values for the estimated *b_1_* coefficients from Equation 1). Of these SNPs, the first 50,000 are clearly those that explain nearly all of the heritability. The line slightly above, labeled GWGEI-G, removes blocks of 5000 SNPs as ranked by lowest p-values for the main effect of the SNP from the GWGEI analysis (the *b_1_* coefficient from Equation 2). The general pattern is roughly similar to removal based on the GWAS analysis although there is still non-trivial heritability remaining after the removal of the first 50,000 and even 100,000 SNPs.

**Figure 1:**
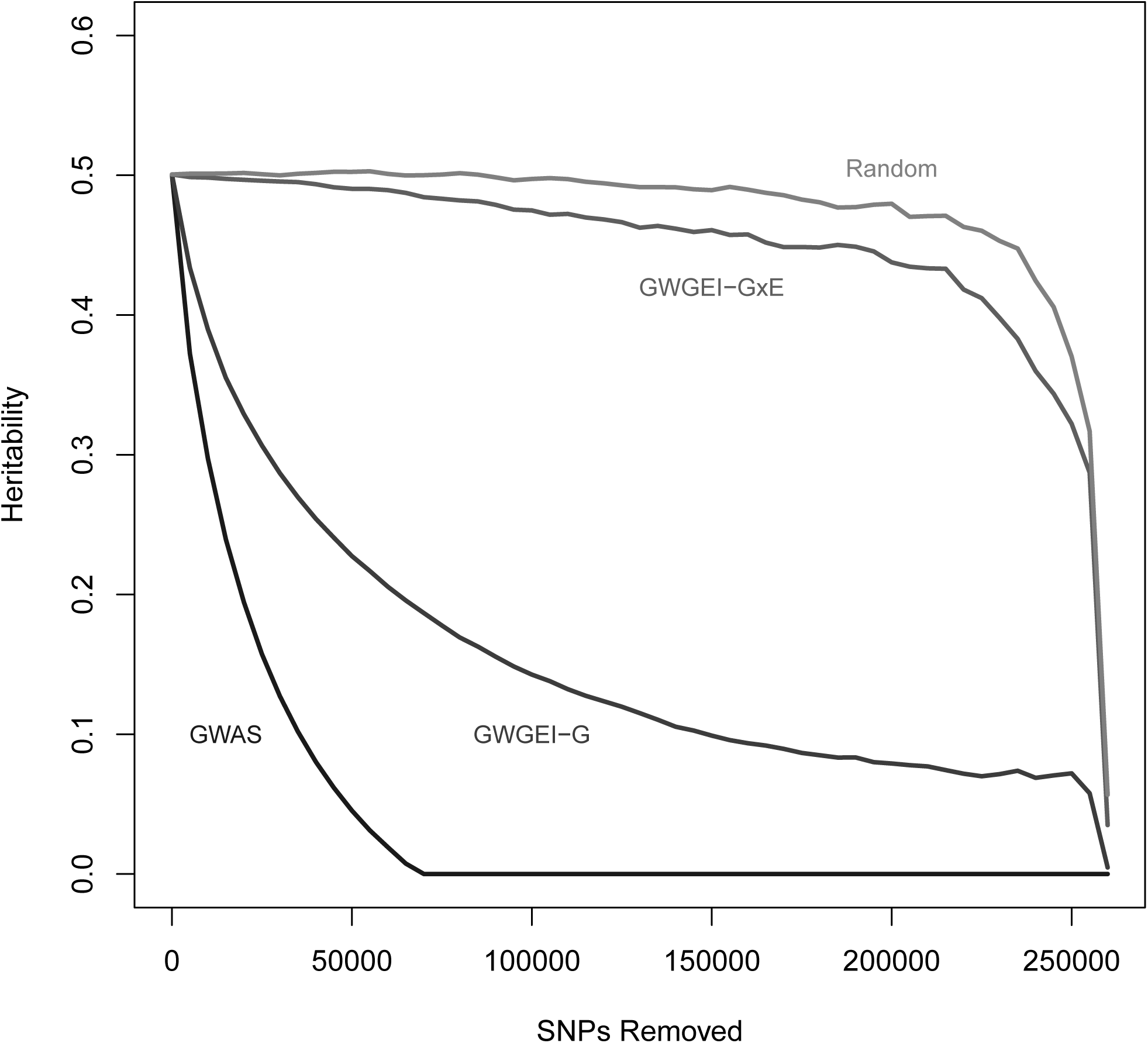
Heritability as a function of remaining SNPs: Removed SNPs for different curves are ordered by p-values from corresponding genome-wide analyses.

The line labeled GWGEI-GxE is the most interesting of the four. This line is based on removing SNPs as ordered by the p-values of the SNP-environment interaction from the GWGEI analysis (the *b*_5_ coefficient from Equation 2). The first 200,000 SNPs, as ordered by this analysis, explain only a fraction of the overall heritability. This curve can be compared to the curve labeled Random which is based on removing SNPs that are randomly ordered. Essentially, the top hits from the GWGEI-GxE SNPs explain no more of the heritability than randomly chosen SNPs.

Figure 2 is based on a separate set of GWAS analyses. The same model as in Equation 3 is used except there is no control for environment (*E_i_*, does not appear). Instead, analyses are run separately on those with and without a college degree. The appropriately labeled curves in Figure 2 correspond to these separate analyses. Note that for both groups, the top hits in the GWAS explain the majority of the heritability using fewer SNPs than in the data as a whole. This can be observed by comparing the No College and College lines to the GWAS line, which is a replication of the line from Figure 1 (note that the scale of the x-axis has changed as well).

**Figure 2:**
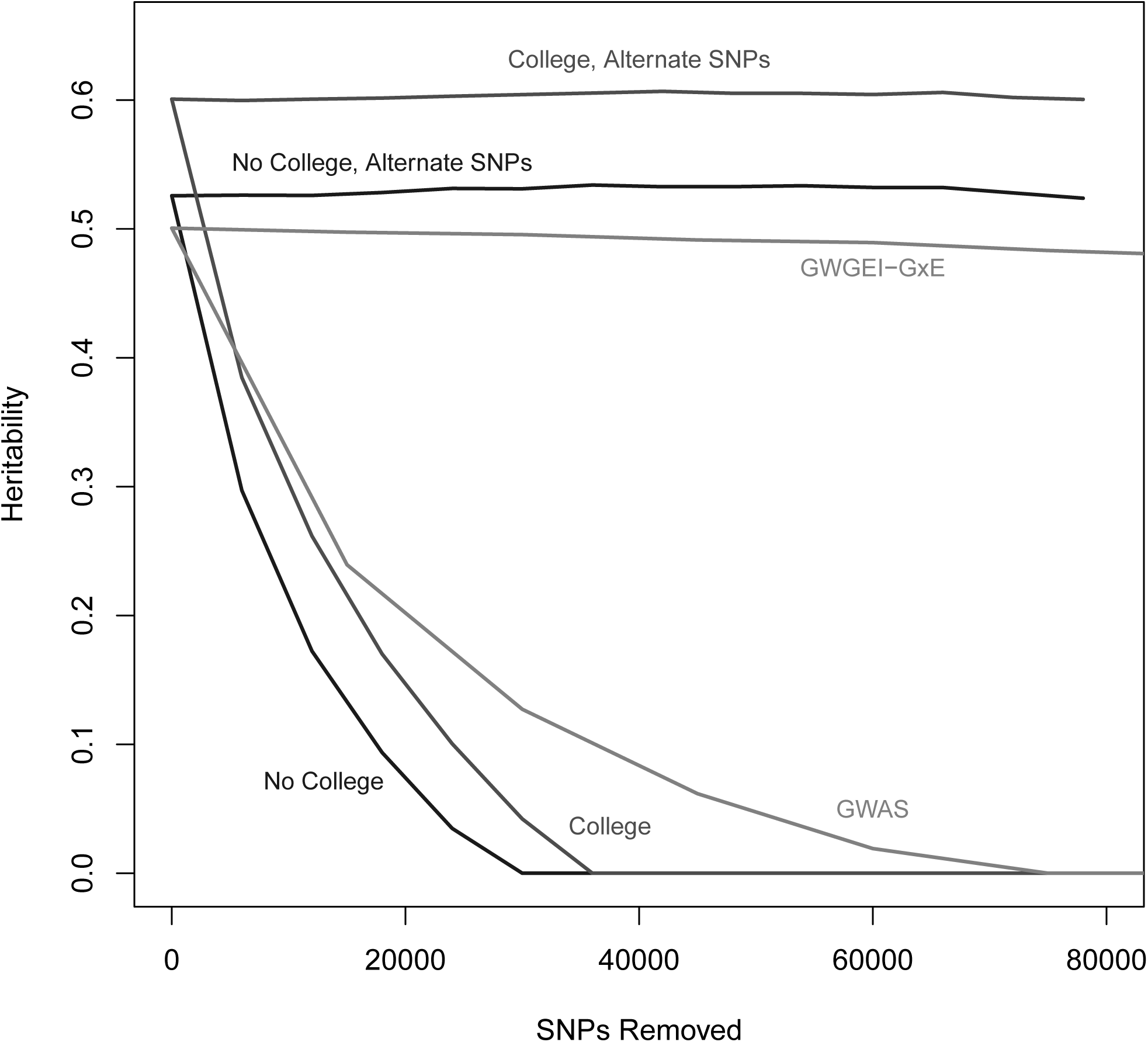
Heritability as a function of remaining SNPs: Removed SNPs for different curves are ordered by p-values from corresponding genome-wide analyses done separately on respondents that had and had not completed college separately.

In contrast, the curves at the top of Figure 2 remove SNPs for the groups as they are ordered from the GWAS in the other group. So, the line labeled No College, Alternate SNPs computes heritability for the group of individuals who did not finish college based on removing SNPs in the order they were ranked from the GWAS using the college graduates. What is remarkable is that essentially *no* heritability is explained by this process. This finding is primarily a reminder that the Framingham sample is underpowered for true identification of causal variants.

### 4.1 GCTA compared to family-based heritabilities

The GCTA approach discussed here utilized genetics data to compute heritability estimates, but the family-based structure of Framingham allows us to compare this estimate to traditional heritability estimates based on family trios (mother, father, child). Using the approach described in Rabe-Hesketh, Skrondal, and Gjeesing (2008) and modern software for estimation of Bayesian models (Stan Development Team, 2012), we obtained a Bayesian estimate for the heritability of BMI. This approach allowed us to also control for age and gender of respondents. These are shown in Figure 3. Bayesian Estimates were centered around 0.355 with a 95% CI from 0.34 to 0.4. A GCTA analysis performed using both waves 2 and 3 (N=3603) led to a heritability of 0.42 (0.03). However, this GCTA estimate is not directly comparable due to the family-based estimate due to the presence of the covariates. An additional GCTA estimate that controlled for age and gender was only slightly lower, 0.41 (0.03). It is reassuring that the two approaches are yielding comparable results and suggest some reasons for why the family-based approach might be computing smaller heritability estimates in the discussion.

**Figure 3:**
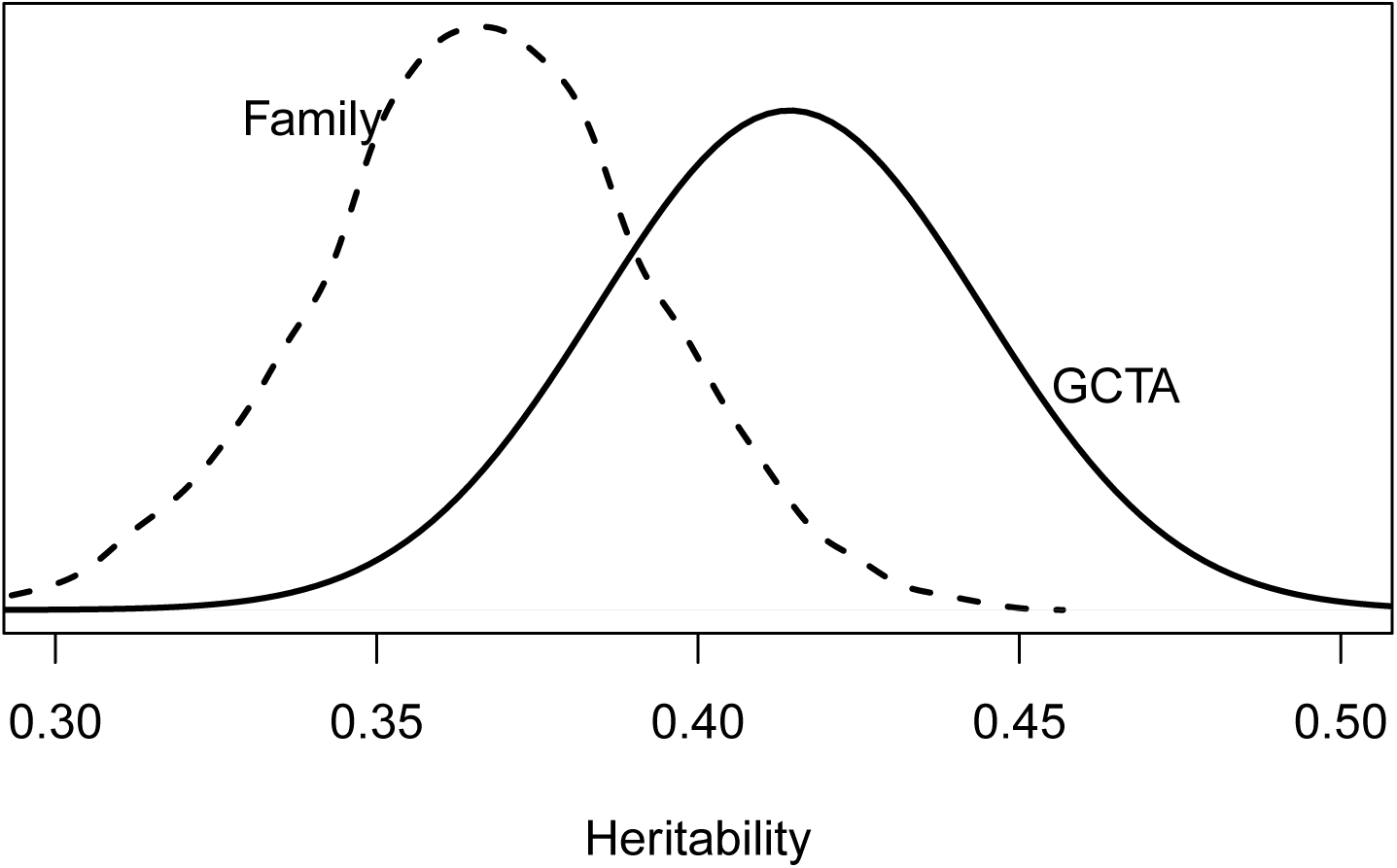
Comparison of GCTA and family-based heritability estimates

## 5 Discussion

This paper focused on the heritability that remained in a group of individuals after removing SNPs in different orders. Figure 1 compared the heritability of BMI in the third generation Framingham after removing SNPs as they were ordered by their p-values in different genome-wide association analyses. Our first main result was a demonstration that the top hits indicated by a traditional GWAS explained nearly all of the heritability. In contrast, those hits indicated by a gene-environment interaction explained essentially none of the heritability. The second main result is shown in Figure 2. The key point is that the top hits from a GWAS using only those that graduated from college explained none of the heritability for those that did not attend college. The opposite version (heritability for those who attended college after removing top hits from a GWAS on those that did not attend college) was also true. Although power^3^ is clearly a concern here, this may also suggest that SNP effect estimates are potentially extremely sensitive to environment.^4^

The third main result was a comparison of heritability estimates based on measured genoy-type versus family structure. The estimates were roughly comparable, which is especially interesting given that the heritability estimates in Figure 3 are based on different sources of information. The family-based estimate uses only indicated familial relationships between waves 2 and 3 while the GCTA estimate is based on measured phenotype. Moreover, the GCTA estimate explicitly excludes pairs with a genetic relationship over some (relatively low) threshold so that family pairs don’t bias the estimate due to the shared environment. The fact that the GCTA estimate of heritability is higher than the family-based estimate is interesting. One potential problem is that the family-based estimate isn’t accounting for the fact that some families have multiple siblings. However, this would probably bias this estimate upwards and so is unlikely to explain the difference.

The results of this research echo those of Boardman et al. (2014) which question the contribution of genetic studies that focus on specific SNPs for social science in general, especially given the limited sample sizes that characterize most of the studies in question. However, we do think that information based on the whole genome could be quite useful. Heritabilities are one approach that can potentially provide information about genetic causes of behavior, but working strictly with heritabilities might be limiting since they don’t allow one to simultaneously control for genetic and environmental factors. A more interesting possibility is the consideration of polygenic scores (e.g., Belsky et al., 2013), which are more tractable in the statistical models commonly used by social scientists.

## Footnotes

1 http://www.complextraitgenomics.com/software/gcta/

2 The pruning of the SNPs was done in the full Framingham sample, not in the subsample of the W3 respondents.

3 Not simply power to detect GWAS effects, but power to detect significant heritabilties via GCTA. See Visscher et al. (2014).

4 It is worth noting that we also found, in Boardman et al. (2014), that SNP effects are measured with sufficient noise that they fail to replicate in a random split of the Framingham data.

## References

Belsky, D. W., Moffitt, T. E., Sugden, K., Williams, B., Houts, R., McCarthy, J., & Caspi, A. (2013). Development and evaluation of a genetic risk score for obesity. Biodemography and social biology, 59(1), 85–100.

Boardman, J. D., Domingue, B. W., Blalock, C. L., Haberstick, B. C., Harris, K. M., & McQueen, M. B. (2014). Is the gene-environment interaction paradigm relevant to genome-wide studies? the case of education and body mass index. Demography, 51(1), 119–139.

Boardman, J. D., Roettger, M. E., Domingue, B. W., McQueen, M. B., Haberstick, B. C., & Harris, K. M. (2012). Gene-environment interactions related to body mass: School policies and social contetx as environmental moderators. Journal of Theoretical Politics, 24(3), 370–388.

Conley, D., Siegal, M. L., Domingue, B. W., Harris, K. M., McQueen, M. B., & Boardman, J. D. (2014). Testing the key assumption of heritability estimates based on genome-wide genetic relatedness. Journal of human genetics, 59(6), 342–345.

Domingue, B. W., Wedow, R., Conley, D., McQueen, M., Hoffman, T., & Boardman, J. (in press). Genome-wide estimates of heritability for social demographic outcomes. Biodemography and Social Biology.

Fox, C., Heard-Costa, N., Cupples, L., Dupuis, J., Vasan, R., & Atwood, L. (2007). Genome-wide association to body mass index and waist circumference: the Framingham heart study 100k project. BMC Medical Genetics, 8(Supplement 1), S18.

Haberstick, B., Lessem, J., McQueen, M., Boardman, J., Hopfer, C., Smolen, A., & Hewitt, J. (2010). Stable genes and changing environments: Body mass index across adolescence and young adulthood. Behavior Genetics, 40, 495–504.

Kumar, S. K., Feldman, M. W., Rehkopf, D. H., & Tuljapurkar, S. (2016). Limitations of gcta as a solution to the missing heritability problem. Proceedings of the National Academy of Sciences, 113(1), E61–E70.

Laird, N., Horvath, S., & Xu, X. (2000). Implementing a unified approach to family based tests of association. Genetic Epidemiology, 19(Supplement 1), S36–S42.

Laird, N., & Lange, C. (2006). Family-based designs in the age of large-scale gene-association studies. Nature Review Genetics, 7, 385–394.

Lee, Y., Lai, C., Ordovas, J., & Parnell, L. (2011). A database of gene-environment interactions pertaining to blood lipid traits, cardiovasculat disease, and type 2 diabetes. Journal of Data Mining in Genomics and Proteonics (1), 1–8.

Loos, R., & Bouchard, C. (2003). Obesity-is it a genetic disorder? Journal of Internal Medicine, 254, 401–425.

Maher, B. (2008). Personal genomes: The case of the missing heritability. Nature, 456, 18–21.

Manolio, T., Collins, F., Cox, N., Goldstein, D., L.A. Hindorff, Hunter, D. Visscher, P. (2010). Finding the missing heritability of complex diseases. Nature, 461(7265), 747–753.

Moreno-Macias, H., Romieu, I., London, S., & Laird, N. (2010). Gene-environment interaction tests for family studies with quantitative phenotypes: a review and extension to longitudinal measures. Human Genomics, 4(5), 302–326.

Rabe-Hesketh, S., Skrondal, A., & Gjeesing, H. (2008). Biometrical modeling of twin and family data using standard mixed model software. Biometrics, 64, 280–288.

Shanahan, M., & Hofer, S. (2005). Social context in gene-environment interactions: Retrospect and prospect. Journals of Gerontology: Series B, 60B, 65–76.

Speed, D., Hemani, G., Johnson, M. R., & Balding, D. J. (2012). Improved heritability estimation from genome-wide snps. The American Journal of Human Genetics, 91(6), 1011–1021.

Splansky, G., Yang, D. C. Q., Atwood, L., Cupples, L., Benjamin, E., D’Agostino, R. Levy, D. (2007). The third generational cohort of the national heart, lung, and blood institute’s Framingham heart study: Design, recruitment, and intial examination. American Journal of Epidemiology, 165(11), 1328–1335.

Stan Development Team. (2012). Stan: A C++ library for probability and sampling [Computer software manual]. Retrieved from http://mc-stan.org/ (Version 1.0)

Visscher, P. M., Hemani, G., Vinkhuyzen, A. A., Chen, G.-B., Lee, S. H., Wray, N. R., Yang, J. (2014). Statistical power to detect genetic (co) variance of complex traits using snp data in unrelated samples. PLoS Genet, 10(4), e1004269.

Yang, J., Benjamin, B., McEvoy, B., Gordon, S., Henders, A., Nyholt, D. Visscher, P. (2010). Common snps explain a large proportion of the heritability for human height. Nature Genetics, 42(7), 565–569.

Yang, J., Lee, S., Goddard, M., & Visscher, P. (2011). GCTA: A tool for genome-wide complex trait analysis. American Journal of Human Genetics, 88(1), 76–82.

Yang, W., Kelly, T., & He, J. (2007). Genetic epidemiology of obesity. Epidemiologic Reviews, 29, 49–61.

